# Liver-specific glucocorticoid action in alcoholic liver disease: study of glucocorticoid receptor knockout and knockin mice

**DOI:** 10.1101/2023.10.04.557166

**Authors:** Yazheng Wang, Hong Lu

## Abstract

**Background:** Glucocorticoids are the only first-line drugs for severe alcoholic hepatitis (AH), with limited efficacy and various side effects on extrahepatic tissues. Liver-targeting glucocorticoid therapy may have multiple advantages over systemic glucocorticoid for AH. The purpose of this study was to determine the role of hepatocellular glucocorticoid receptor (GR) in alcoholic steatosis (AS) and AH.

**Methods:** AS was induced by a high-fat diet plus binge alcohol in adult male and female mice with liver-specific knockout (LKO) and heterozygote of GR. AH was induced by chronic-plus-binge in middle-aged male mice with liver-specific knockin of GR. Changes in hepatic mRNA and protein expression were determined by qPCR and Western blot.

**Results:** GR LKO aggravated steatosis and decreased hepatic expression and circulating levels of albumin in both genders of AS mice but only increased markers of liver injury in male AS mice. Marked steatosis in GR LKO mice was associated with induction of lipogenic genes and down-regulation of bile acid synthetic genes. Hepatic protein levels of GR, hepatocyte nuclear factor 4α, and phosphorylated STAT3 were gene-dosage-dependently decreased, whereas that of lipogenic ATP citrate lyase was increased in male GR heterozygote and LKO mice. Interestingly, hepatic expression of estrogen receptor α (ERα) was induced, and the essential estrogen-inactivating enzyme sulfotransferase 1E1 was gene-dosage-dependently down-regulated in GR heterozygote and knockout AS mice, which was associated with induction of ERα-target genes. Liver-specific knockin of GR protected against liver injury and steatohepatitis in middle-aged AH mice.

**Conclusions:** Hepatocellular GR is important for protection against AS and AH.

## INTRODUCTION

According to the World Health Organization: there are about 15-20 million people worldwide who abuse alcohol, of which 10-20% (1.5 to 4 million) have varying degrees of alcoholic liver disease (ALD). Statistics show that 40% to 90% of the 26,000 people who die from cirrhosis each year are associated with ALD (Organization et al., 2018). Alcohol abuse can lead to three kinds of liver injury, and they usually develop in the following order: fatty liver or hepatic steatosis, alcoholic hepatitis (AH), and finally, cirrhosis. AH patients have severe inflammation and cholestatic liver injury (Savolainen et al., 1995), and severe AH (sAH) has a high mortality rate (Stepanova et al., 2010). Glucocorticoids (GCs) are currently the only available drug therapy for sAH (Dugum and McCullough, 2016), however, prednisolone, the current standard of care for sAH, only marginally reduces 28-day mortality without long-term improvement (Thursz et al., 2015). Dissecting the beneficial and detrimental actions of GC therapy in sAH is a significant challenge in basic research, drug development, and personalized medicine.

Marked steatohepatitis, jaundice, cholestatic liver injury, and impaired liver regeneration are hallmarks of sAH which causes high mortality (Sidhu et al., 2017, Hosseini et al., 2019). Literature suggests hepatic GR deficiency in fatty liver and AH (Ahmed et al., 2012, Nasiri et al., 2015, Lu, 2022). Most recent literature support a protective role of hepatocellular GR against steatosis during high fatty acid influx (Nasiri et al., 2015, Roqueta-Rivera et al., 2016, Quagliarini et al., 2019, Lu et al., 2022, Mueller et al., 2011). The marked decrease of urea synthesis in AH is associated with the disease severity and hepatic encephalopathy (Glavind et al., 2016, Parekh and Balart, 2015). GR controls the urea cycle in the liver (Okun et al., 2015), and prednisolone restores urea synthesis in survivors of sAH (Glavind et al., 2017). Additionally, cholestasis correlates highly with malnutrition and AH severity (Trinchet et al., 1994, Alpert and Hart, 2016, Axley et al., 2017, Nissenbaum et al., 1990). GR in hepatocytes is anti-apoptotic and anti-inflammatory (Mueller et al., 2011, Gruver-Yates and Cidlowski, 2013), and activation of GR protects against cholestatic liver injury and steatohepatitis in patients (Zollner and Trauner, 2009) and mice (Petrescu et al., 2017). All these hepatoprotective and anti-inflammatory effects of GR activation on hepatocytes likely contribute to the beneficial effects of current GC therapy in sAH. Additionally, hepatic GR is essential in protecting against liver failure and mortality in sepsis (Jenniskens et al., 2018), a leading cause of death of sAH (Gustot et al., 2017, Perez-Hernandez et al., 2017).

Long-term treatment with GCs can cause a series of adverse effects and is the main reason for limiting AH treatment, the severity of which is proportional to the dose and duration of GC administration (Huscher et al., 2009). GR activation in extrahepatic tissues promotes alcohol consumption and psychiatric problems (Jimenez et al., 2020, Vendruscolo et al., 2015), adipose lipolysis (Shen et al., 2017), intestinal bile acid (BA) reabsorption and cholestasis, gut dysbiosis, and gastrointestinal bleeding (Narum et al., 2014, Out et al., 2014, Shukla et al., 2022), and skeletal muscle wasting (Lang et al., 2007). Importantly, high doses of GC can exacerbate alcoholic liver injury and delays liver regeneration by inhibiting macrophage-mediated phagocytic and hepatic regenerative functions (Kwon et al., 2014, Kim et al., 2016, Wynn and Vannella, 2016, Gao and Shah, 2015). In contrast, GR deficiency in hepatocytes impairs liver regeneration (Shteyer et al., 2004). Therefore, liver-specific delivery of GCs should markedly improve GC therapy of sAH by ameliorating the adverse effects of GCs on extrahepatic tissues and immunosuppression.

Estrogen (E2) signaling is mediated primarily via the nuclear hormone receptors estrogen receptor alpha (ERα) and ERβ. In female mammals, ERα is highly expressed in the liver, where it acts as a sensor of the nutritional status and adapts the metabolism to the reproductive needs. Impaired ERα function is associated with obesity and metabolic dysfunction in humans and rodents (Rahman and Cao, 2016). E2 induces ERα in periportal hepatocytes, ameliorates liver injury, and promotes liver regeneration (Zhu et al., 2013, Zhu et al., 2018a, Della Torre et al., 2011, Lee et al., 2019, Uebi et al., 2015, Tsugawa et al., 2019, Dubuquoy et al., 2015). Acute binge alcohol studies indicate a female protection from acute AH (Gao et al., 2017, Spruiell et al., 2015, Li et al., 2017). The ERα-target gene growth differentiation factor 15 (GDF15) maintains mitochondrial function and protects against ALD. GR can inhibit the transcriptional activities of ERα via physical interaction with ERα and induction of the essential E2-inactivating enzyme sulfotransferase 1E1 (SULT1E1) (Gong et al., 2008, Karmakar et al., 2013, West et al., 2016, Bolt et al., 2013).

Although GCs are the only first-line drugs recommended for the treatment of sAH (Hosseini et al., 2019), the role of hepatocellular GR in AH has not yet been studied. Mice with the whole-body knockout of GR are embryonic lethal. Knockout of GR specifically in perinatal hepatocytes, using the Alfp-cre, results in the death of ∼50% of the knockout mice within 48 h after birth, and the surviving knockout mice have postnatal growth retardation (Tronche et al., 2004, Opherk et al., 2004). GR plays a key role in early-life programming. These ∼50% survived knockout mice (Tronche et al., 2004, Opherk et al., 2004) most likely already undergo marked genetic reprogramming to survive the perinatal loss of GR, and thus results from the study of conventional liver-specific GR knockout mice may obscure the true role of GR in adult liver.

Overeating high-fat-diet (HFD) aggravates AH and drives nonalcoholic fatty liver disease (NAFLD) (Gao et al., 2017). Literature suggests hepatic GR deficiency in NAFLD and AH (Ahmed et al., 2012, Nasiri et al., 2015, Lu, 2022). The purpose of this study was to use our novel model of mice with tamoxifen-inducible liver-specific knockout (LKO) of GR to determine the effects of hepatic deficiency of GR on alcoholic steatosis (AS) induced by high-fat diet (HFD) feeding plus binge alcohol and the alterations of GR- and ERα-target genes in AS mice with GR deficiency. Additionally, we studied the effect of hepatocyte-specific activation of GR on AH induced by chronic plus binge in male middle-aged mice with liver-specific knockin (LKI) of GR. Results from these studies demonstrate a critical role of hepatocellular GR in the protection against alcoholic liver disease in mice and a marked dysregulation of GR- and ERα-signaling in patients with sAH.

## MATERIALS AND METHODS

### Animals and treatments

#### Mice with liver-specific knockout and heterozygote of GR

We generated the mice with inducible LKO of GR by crossing the GR floxed mice (#021021, Jackson Laboratory) with the SA^+/CreERT2^ mice, which carry a tamoxifen-inducible ERα-fused Cre under the control of an albumin promoter (Schuler et al., 2004, Bonzo et al., 2012). Adult male and female wildtype (WT) (Cre/-), GR liver-specific heterozygote (LHet) (GR fl/+, Cre/+), and GR LKO (GR fl/fl, Cre/+) mice (in C57BL/6 background) were injected tamoxifen 5 mg/kg once daily for 2 d to activate the Cre-ERT2. Ten days after tamoxifen injection, all mice were fed the HFD (#D12492, Research Diets) for 3 weeks. Three weeks after HFD feeding, mice had intragastric administration of ethanol 5 g/kg (50% 10 ml/kg, AH groups) or isocaloric maltose (HFD groups) 9 g/kg (0.45 g/ml 20 ml/kg) in the morning. All mice were sacrificed 9 h after ethanol treatment to collect blood and tissues (Chang et al., 2015). Liver tissues were snap-frozen in liquid nitrogen upon collection and stored at −80 °C until use. Mouse blood samples were taken by orbital bleeding and then centrifuged at 8000 rpm for 10 min. The resultant supernatants were collected as serum samples and stored at −80 °C until future use.

#### Mice with liver-specific knockin (LKI) of GR

We generated PiggyBac (Ding et al., 2005) GR knockin (GR-KI) vector (VectorBuilder Inc.), in which the loxP-flanked STOP cassette prevents the strong CAG-promoter-driven expression of human GRα mRNA (NM_001018074). We bred the PiggyBac GR-KI founder mice (Cyagen Inc.) with SA^+/CreERT2^ mice to generate GR LKI mice, and verified hepatic functional expression of hGR. We determined effect of GR LKI on chronic-plus-binge (E10d+1B)-induced AH (Bertola et al., 2013b) in middle-aged (10-12-month-old) male GR LKI (GR-KI +, Cre/+) and age-matched WT littermates (Cre/-) one month after tamoxifen injection to cause liver-specific expression of the human GR transgene. Briefly, mice were given free access to the chronic drinking (6.7% ethanol in liquid diet, F1258SP, Bio-Serv) for 10 days, followed by one binge alcohol (5g/kg) in the morning. Control GR LKI and WT mice were fed the control liquid diet (F1259SP, Bio-Serv) followed by isocaloric maltose 9 g/kg (0.45 g/ml 20 ml/kg). All mice were sacrificed 9 h after ethanol treatment to collect blood and tissues for analysis. Liver tissues were snap-frozen in liquid nitrogen upon collection and stored at −80 °C until use. Mouse blood samples were taken by orbital bleeding and then centrifuged at 8000 rpm for 10 min. The resultant supernatants were collected as serum samples and stored at −80 °C until future use.

### Determination of blood chemistry as well as lipids in mouse liver samples

Lipids of frozen mouse liver samples were extracted with chloroform: methanol (2:1) and concentrated by vacuum(Lu et al., 2011). The lipid pellets were dissolved in a mixture of 270 μl of isopropanol and 30 μl of Triton X-100. Triglycerides (TG) and total cholesterol (CHO) in the liver and serum were determined using commercial triglyceride and cholesterol analytical kits with standards (Pointe Scientific, Canton, MI). Serum alanine aminotransferase (ALT), bilirubin, and albumin were determined using commercial ALT, bilirubin, and albumin analytical kits with standards (Pointe Scientific, Canton, MI).

### Quantification of serum corticosterone by ELISA

The DetectX® Corticosterone Enzyme Immunoassay Kit (K014-H1, Arbor Assays LLC, Ann Arbor. MI, USA) was used to determine serum corticosterone levels. Serum samples were prepared and analyzed following the manufacturer’s protocol. The online tools from MyAssays provided by Arbor Assays LLC were used to calculate the data.

### RNA isolation and real-time PCR quantification of mRNA

Total RNA from liver tissues was extracted using RNA STAT-60 (Tel-Test, Friendswood, TX, USA) according to the manufacturer’s instructions. Equal amount of total RNAs from each sample in the given group was mixed to prepare the pooled RNA samples. cDNA was produced using a cDNA Synthesis kit (iScript™ cDNA Synthesis Kit, Bio-Rad, Hercules, CA, USA). The diluted cDNA was used for real-time PCR quantification of mRNA using iTaq SYBR® Green Supermix (Bio-Rad, Hercules, CA, USA) and CFX Real-Time PCR Detection System (Bio-Rad). The data were analyzed by CFX Maestro qPCR Analysis Software (Bio-Rad), and the amounts of mRNA were calculated using the CQ value, normalized to an endogenous reference, phosphoglycerate kinase 1 (Pgk1) (Panina et al., 2018). The primers for PCR are listed in Supplemental Table 1.

### Western blot quantification of liver proteins

Briefly, the mouse liver was homogenized in hypotonic buffer (20 mM HEPES-KOH (pH 7.8), 5 mM potassium acetate, 0.5 mM MgCl_2_, 1X protease and phosphatase inhibitor). After centrifugation at 1500 g for 5 min (4 °C), the supernatant was removed (as cytosol), and the rest of the supernatant and most of the upper lipid layer were removed and discarded. The cytosol was centrifuged again 14000 g at 4 °C for 5 min to remove additional lipid/nucleus. The nuclei pellets were washed with hypotonic buffer and centrifuged again at 1500 g, and the resultant pelleted nuclei were incubated at 4°C for 1 h in RIPA buffer (10 mM Tris-HCL (pH 8.0),1 mM EDTA, 0.5 mM EGTA, 1% Triton X-100, 0.1% sodium deoxycholate, 0.1% SDS, 140 mM NaCl and phosphatase inhibitor). After centrifugation of the nuclear/RIPA samples at maximum speed for 10 min (4 °C), the supernatants were collected as nuclear protein samples.

Equal amount of total nuclear or cytosolic proteins from each sample in the given group was mixed to prepare the pooled protein samples for Western blot. Western blot quantification of ATP citrate lyase (ACLY), 70-kDa heat shock protein (HSP70), phosphorylated eukaryotic translation initiation factor 4E (eIF4E)-binding protein 1 (4E-BP1) (P-4EBP1), HSP90, extracellular signal-regulated kinase 1/2 (ERK1/2), ERα, and GR in liver nuclear and cytosol extracts was carried out with the primary antibodies as follows: ACLY (D1X6P) (#13390), HSP70 (#4872), p-4E-BP1 (Thr37/46) (236B4) (#2855), HSP90 (C45G5) (#4877), Phospho-p44/42 MAPK (Erk1/2) (Thr202/Tyr204) (D13.14.4E) XP® (#4370), Phospho-(Tyr705) signal transducer and activator of transcription 3 (pSTAT3, #9145T), phospho-mTOR (Ser2448) (#5536), phospho-mTOR (Ser2481) (#2974), and GR (D6H2L) XP® (#12041) from Cell Signaling Technologies (Danvers, MA, USA). The ERα rabbit pAb (A0296) was purchased from Abclonal (Wobum, MA). Proteins in cytosolic or nuclear extracts from mouse livers were resolved in sodium dodecyl sulphate-polyacrylamide gel electrophoresis using the mini-PROTEAN TGX Stain-Free Precast Gels from Bio-Rad. Primary antibodies were revealed with HRP-conjugated secondary antibodies (anti-rabbit IgG #7074, Cell Signaling) and ECL Western Blotting Substrate (1705061, Bio-Rad). Image Lab software (Bio-Rad) was used to capture signals and determine signal intensities, and the target proteins were normalized to the loaded total proteins in each lane.

### Statistical analysis

Data are presented as mean ± standard error (SE) (pooled samples for female HFD groups in real-time PCR). The student’s t-test was used to determine the statistical difference between WT and KO group, HFD and AS group, as well as the Male and Female group (Prism 8). Statistical significance was set at * p < 0.05 vs. corresponding WT group; # p < 0.05 vs. corresponding HFD group; $ p<0.05 vs. corresponding male group.

## RESULTS

### GR LKO mice had more liver injury than WT mice in AS

We attempted to explore the hepatoprotective mechanisms of GR and the adaptive changes in GR LKO mice. Data showed that LKO of GR increased liver/body weight ratio, and GR-LKO males had more increase in the liver/body weight ratio than GR-LKO females (Fig. 1A). Both male and female HFD-fed GR-LKO mice had elevated hepatic triglycerides (TG) (Fig. 1B) and cholesterol (Fig. 1C) compared to the HFD-fed WT mice. Thus, hepatic GR deficiency led to the accumulation of TG and cholesterol in the liver, indicating an important role of hepatic GR in protecting against HFD-induced NAFLD. Ethanol treatment caused marked increases in hepatic TG and cholesterol only in WT and GR-LKO male mice but not in females which had higher basal hepatic lipids when fed the HFD but were resistant to the ethanol-induced increase of hepatic lipids and liver injury.

**Figure 1.**
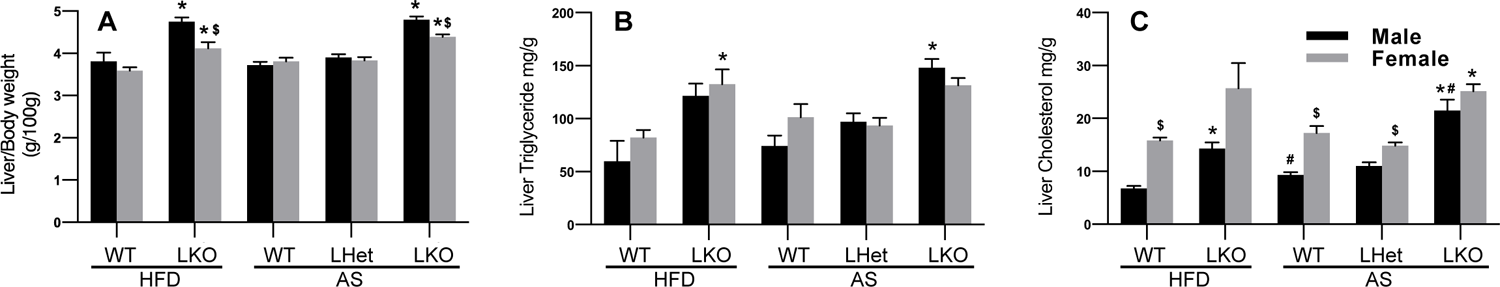
Liver/body weight ratio (A), hepatic triglycerides (B), and hepatic cholesterol (C) in male and female wildtype (WT), GR liver-specific heterozygous (LHet) and GR liver-specific knockout (LKO) HFD-fed and alcoholic steatosis (AS) mice. Mice were fed a HFD for 3 weeks followed by ig administration of ethanol 5 g/kg (AS groups) or isocaloric maltose 9 g/kg (HFD groups) in the morning. All mice were sacrificed 9 h after ethanol treatment to collect blood and tissues for analysis. N=3-7 per group, mean ± SE. * p < 0.05 vs corresponding WT group; # p < 0.05 vs corresponding HFD group; $ p < 0.05 vs corresponding male group.

Histology analysis (H&E staining) found mainly microvesicular steatosis in WT AS mice (Fig. 2C, 2G), whereas massive increases in macrovesicular and microvesicular steatosis were found in both the HFD-fed (Fig. 2B, 2F) and AS (Fig. 2D, 2H) GR-LKO males and females compared to the corresponding WT males (Fig. 2A, 2C) and females (Fig. 2E, 2G). There was no marked difference in histology between the male and female GR-LKO AS mice.

**Figure 2.**
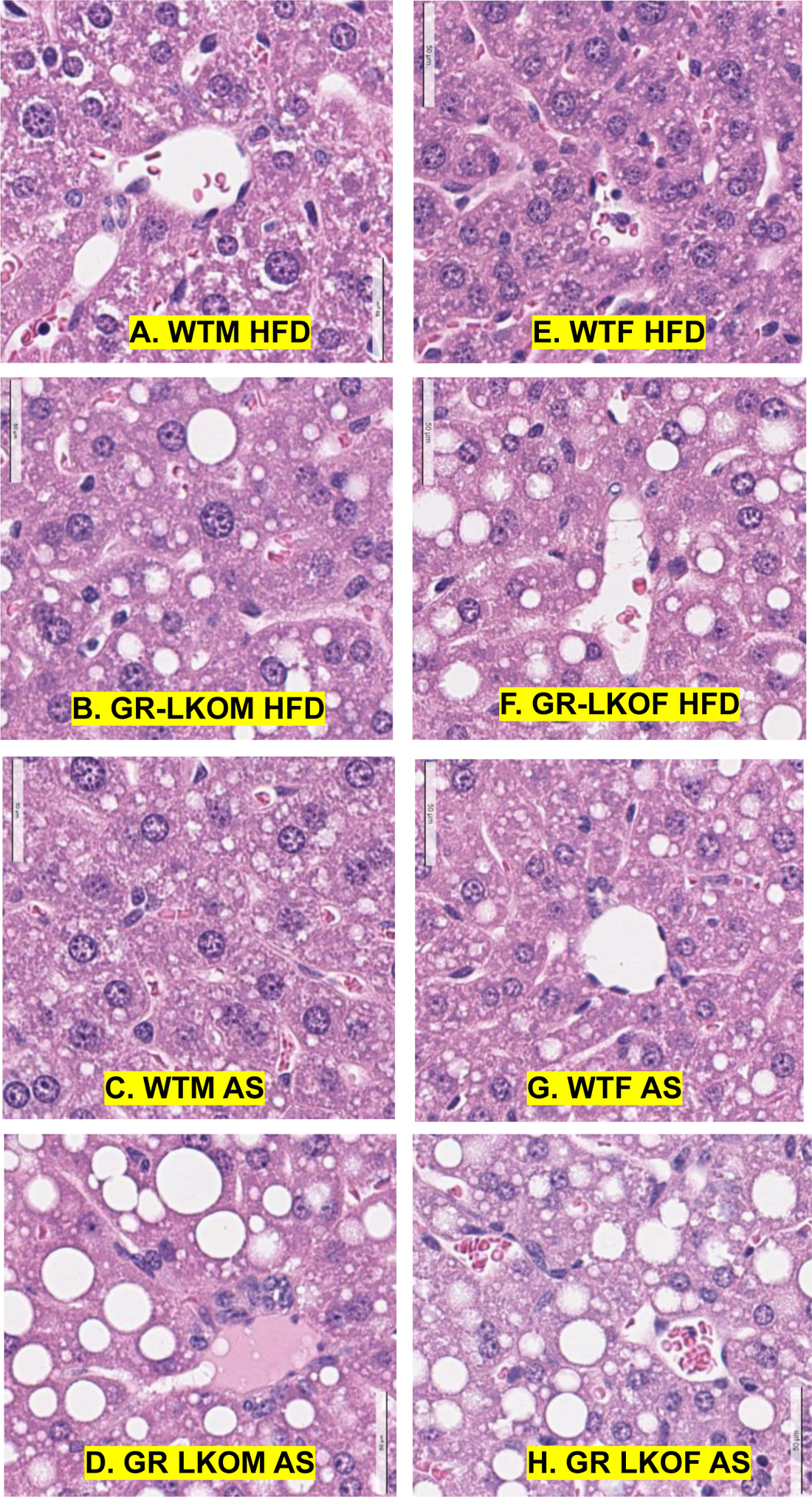
Histology (H & E staining) of livers in wildtype (WT) and GR liver-specific knockout (LKO) male (left) and female (right) HFD-fed and alcoholic steatosis (AS) mice. Mice were fed a HFD for 3 weeks followed by ig administration of ethanol 5 g/kg (AS groups) or isocaloric maltose 9 g/kg (HFD groups) in the morning. All mice were sacrificed 9 h after ethanol treatment to collect blood and tissues for analysis.

Consistent with literature (Gao et al., 2017, Spruiell et al., 2015, Li et al., 2017), we found less liver injury in female AS mice than in male AS mice, manifested by the lack of elevation of blood levels of ALT in WT and GR-LKO AS female mice (Fig. 3A). Compared to WT HFD-fed mice, ALT tended to be increased in ethanol-treated WT males (p = 0.08) and was significantly elevated in GR-LHet and GR-LKO AS male mice. The serum bilirubin levels were not altered by GR deficiency or ethanol treatment, although the bilirubin levels were higher in male mice than in female mice (Fig. 3B). Regarding hepatocellular synthetic function, serum albumin levels were significantly lower in alcohol-treated AS mice (Fig. 3C). Interestingly, serum albumin levels were diminished in both the GR-LHet and GR-LKO mice compared to WT mice, indicating a critical role of hepatic GR in maintaining blood levels of albumin (Fig. 3C). Blood levels of TG were similar in HFD-fed male and female WT mice, whereas blood TG was increased in the AS WT females but tended to be decreased in the AS WT males, resulting in 1.5-fold higher blood TG in the AS WT females than the AS WT males (Fig. 3D). These suggest that a shift of TG distribution from blood to the liver may contribute to the steatosis in the AS WT males. In contrast to the higher blood TG in the females than the males, blood cholesterol levels were lower in the females than in the males (Fig. 3E). Besides, serum cholesterol levels were reduced in both ethanol-treated WT and GR-LKO mice, which may contribute to the increases of hepatic lipids in these mice. Similar to a previous study (Mueller et al., 2011), circulating corticosteroids markedly increased in the HFD-fed GR LKO mice (Fig. 3F). Alcohol treatment had no effect on circulating corticosteroids in WT mice but decreased circulating corticosteroids in the GR LKO females (Fig. 3F).

**Figure 3.**
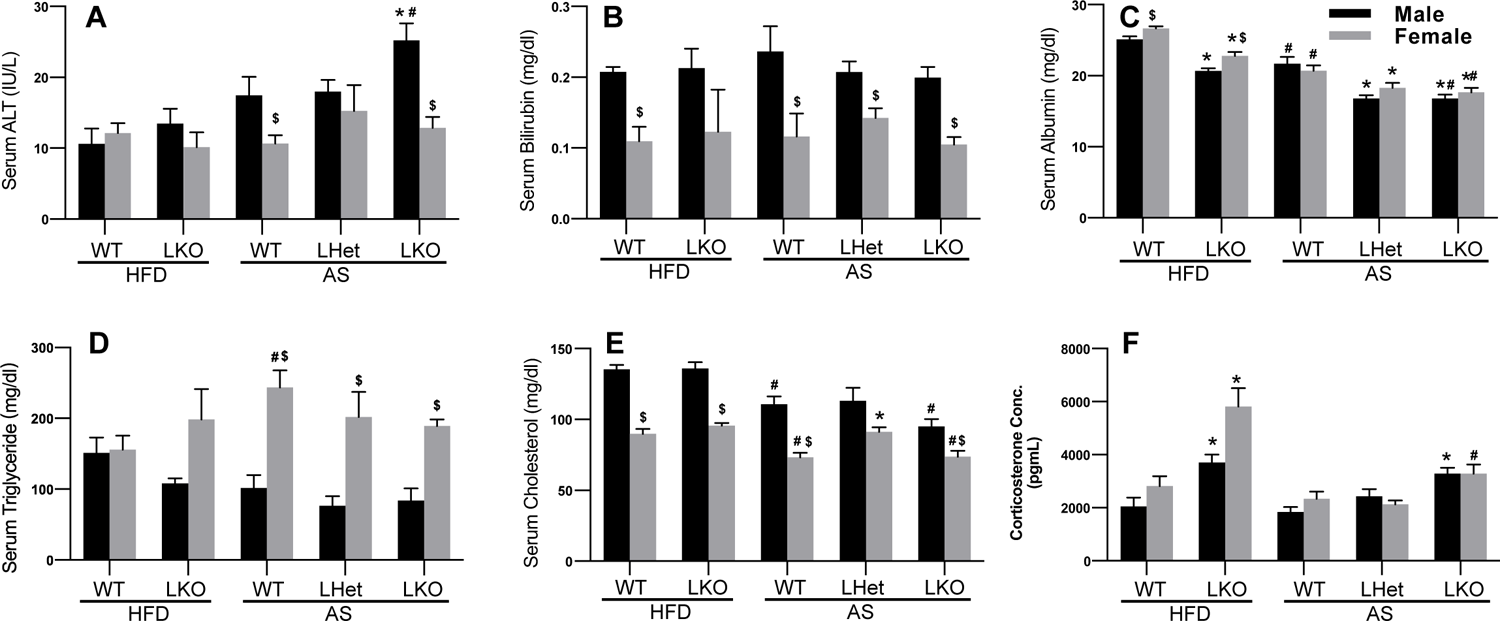
Serum ALT (A), bilirubin (B), albumin (C), triglyceride (D), cholesterol (E), and corticosteroid (F) in male and female wildtype (WT), GR liver-specific heterozygous (LHet) and GR liver-specific knockout (LKO) HFD-fed and alcoholic steatosis (AS) mice. Mice were fed a HFD for 3 weeks followed by ig administration of ethanol 5 g/kg (AS groups) or isocaloric maltose 9 g/kg (HFD groups) in the morning. All mice were sacrificed 9 h after ethanol treatment to collect blood and tissues for analysis. N=3-7 per group, mean ± SE. * p < 0.05 vs corresponding WT group; # p < 0.05 vs corresponding HFD group; $ p < 0.05 vs corresponding male group.

### Changes in hepatic mRNA expression in GR LKO mice

The present study clearly demonstrates a crucial role of hepatic GR in the prevention of fatty liver induced by HFD and alcohol binge. However, we found only moderately increased liver injury in GR LKO AS mice compared to WT AS mice. Some adaptive protective changes may help ameliorate liver injury in the GR LKO AS mice. Thus, we determined changes in hepatic mRNA expression in these mice using RNA-sequencing (RNA-seq) of pooled samples, followed by qPCR verification with individual samples.

Albumin (ALB) is exclusively produced by the liver and low serum albumin levels is associated with liver dysfunction and liver failure (Garcia-Martinez et al., 2013). Consistent with the serum albumin data, hepatic Alb mRNA expression was higher in females than males (Fig. 4A). Alb expression was decreased in both the GR LHet and LKO mice, indicating a critical role of hepatic GR in maintaining hepatic Alb expression and production (Fig. 4A). Hepatic GR deficiency caused gene-dosage-dependent down-regulation of a group of key GR-target cytoprotective genes such as 6-Phosphofructo-2-Kinase/Fructose-2,6-Biphosphatase 3 (Pfkfb3) (Fig. 4B) and metallothionein 1 (Mt1) (Fig. 4C). The rate-limiting glycolytic enzyme PFKFB3 activates the AMP kinase to promote glycolysis and cell survival and inhibit lipogenesis (Wu et al., 2005, Domenech et al., 2015). Metallothionein protects against HFD-induced fatty liver (Sato et al., 2010). Mt1 and Mt2 are the most markedly down-regulated genes in the HFD-fed obesity-prone C57/BL6 mice (Waller-Evans et al., 2013). Hepatic expression of the bile-acid synthetic genes cytochrome P450 7b1 (Cyp7b1) and Cyp39a1 were male- and female-predominant, respectively. Cyp7b1 attenuates NAFLD in mice at thermoneutrality (Evangelakos et al., 2021). In addition to BA synthesis, CYP39A1 inhibits the transcriptional activity of c-MYC via its C-terminal region (Ji et al., 2022). Cyp39a1 was highly gene-dosage-dependently down-regulated in both the male and female GR LHet and LKO AS mice (Fig. 4D), whereas Cyp7b1 was down-regulated in the male, but not female GR LHet and LKO AS mice (Fig. 4E). Activation of sterol regulatory element-binding protein 1 (SREBP-1) enhances lipid synthesis via induction of critical lipogenic enzymes such as stearoyl-CoA desaturase (SCD1) and ATP citrate synthase (ACLY). Unsaturated FAs potently down-regulate SREBP-1C to feedback inhibit lipogenesis (Hannah et al., 2001). Consistently, hepatic Srebp-1c was down-regulated in the WT AS mice; however, such feedback down-regulation of Srebp-1c was impaired in the GR LHet and LKO AS mice. The induction of Srebp-1c in the GR LHet and LKO AS mice (Fig. 4F) was associated with marked induction of the key lipogenic enzymes Scd1 (Fig. 4G) and Acly (Fig. 4H) in these mice. Lipin-1 (Lpin1) deficiency in mice causes lipodystrophy, characterized by impaired adipose tissue development and insulin resistance, and GC is the stimulator of Lpin1 expression in adipocyte differentiation (Zhang et al., 2008). GR binds to the glucocorticoid response element (GRE) in the Lpin1 promoter and leads to transcriptional activation of Lpin1 in hepatocytes and adipocytes (Zhang et al., 2008). Liver-specific Lpin1 deficiency exacerbates experimental alcohol-induced steatohepatitis in mice (Hu et al., 2013). We found a gene-dosage-dependent down-regulation of Lpin1 in GR LHet and LKO AS mice (Fig. 4I).

**Figure 4.**
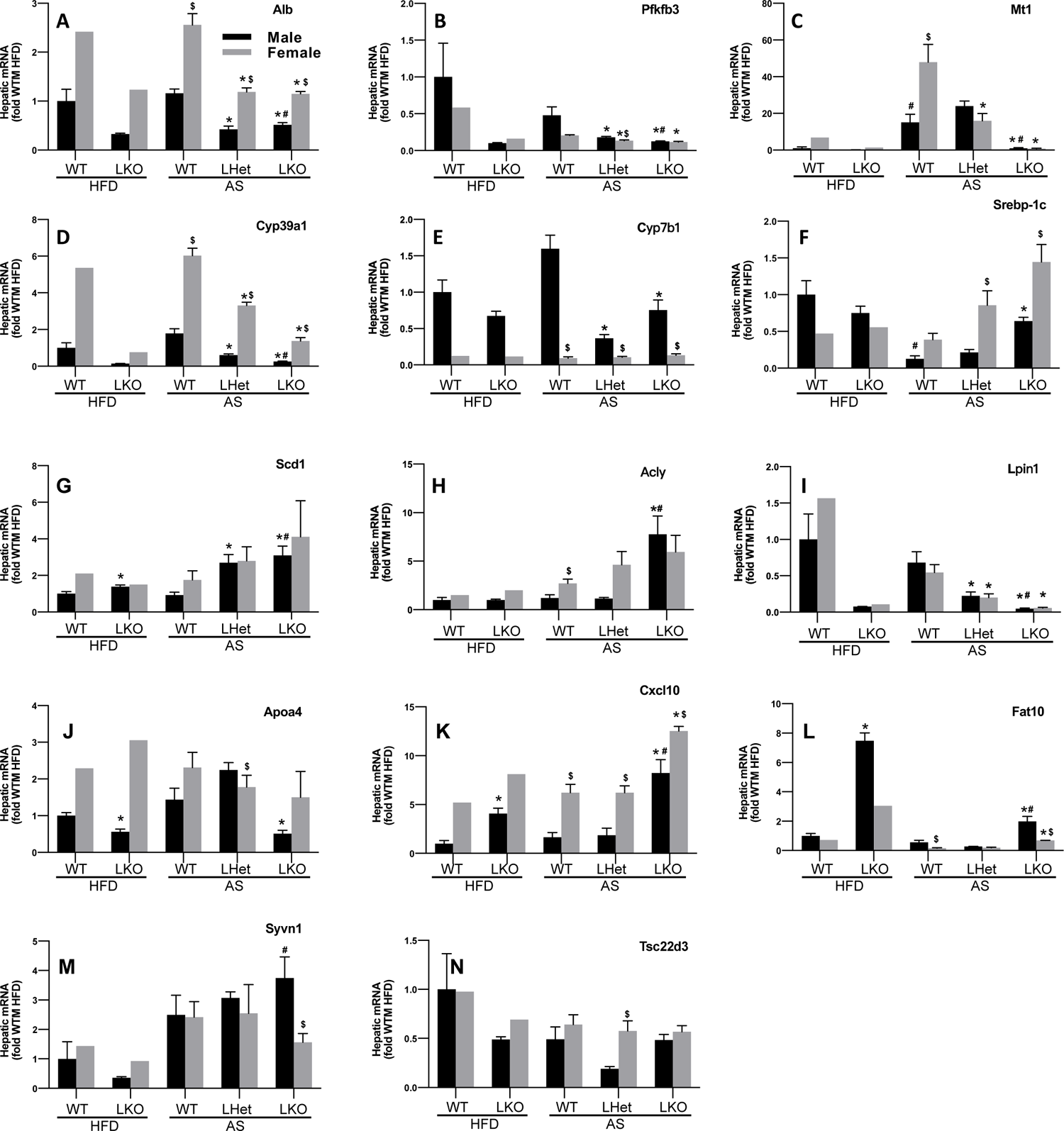
Hepatic mRNA expression in male and female wildtype (WT), GR liver-specific heterozygous (LHet) and GR liver-specific knockout (LKO) HFD-fed and alcoholic steatosis (AS) mice. Mice were fed a HFD for 3 weeks followed by ig administration of ethanol 5 g/kg (AS groups) or isocaloric maltose 9 g/kg (HFD groups) in the morning. All mice were sacrificed 9 h after ethanol treatment to collect blood and tissues for analysis. N=3-7 per group, mean ± SE (pooled samples for WT-HFD and KO-HFD females). * p < 0.05 vs corresponding WT group; # p < 0.05 vs corresponding HFD group; $ p < 0.05 vs corresponding male group.

The apolipoprotein Apo-AIV (Apoa4) promotes hepatic secretion and the expansion of larger very-low-density lipoproteins that are thought to have low cardiovascular risk (VerHague et al., 2013). WT female mice had higher Apoa4 mRNAs than the WT males (Fig. 4J). Hepatic Apoa4 mRNA was down-regulated in GR-LKO AS males, but little changed in GR-LKO AS females, which may contribute to the lower blood levels of TG but more increases in hepatic lipids in GR-LKO AS males than GR-LKO AS females (Fig. 4J).

GC can inhibit the expression of pro-inflammatory factors, such as C-X-C motif chemokine ligand 10 (Cxcl10), a small chemokine (Newton et al., 2017). WT females had higher hepatic Cxcl10 expression than the WT males (Fig. 4K). Cxcl10 was strongly induced in the HFD-fed and AS GR-LKO males but only moderately increased in the GR-LKO AS females (Fig. 4K).

HLA-F-adjacent transcript 10 (FAT10/UBD), a ubiquitin-like modifier, is implicated as a major contributor to AH (Jia et al., 2020). FAT10 mediates the ubiquitin-independent proteasomal degradation and NF-kB activation, and FAT10 is markedly induced in AH and diverse liver diseases (Jia et al., 2020). Fat10 has an important role in the regulation of lipid metabolism, menefisted by the marked decreased adipogenesis in the Fat10 knockout mice (Canaan et al., 2014). We found strong induction of Fat10 in HFD-fed GR-LKO mice (Fig. 4L). Acute alcohol treatment down-regulates FAT10 in hepatocytes (Ganesan et al., 2015). Likewise, ethanol treatment attenuated hepatic induction of Fat10 in GR-LKO mice. Interestingly, Fat10 expression was much higher in AS males than females, suggesting a role of estrogen in preventing hepatic induction of Fat10. In addition to Fat10, synoviolin 1 (SYVN1) is an E3 ligase that promotes adipogenesis (Fujita et al., 2015). Moreover, SYVN1 suppresses nuclear factor-erythroid factor 2-related factor 2 (Nrf2)-mediated cellular protection during liver cirrhosis via enhanced Nrf2 ubiquitylation and degradation (Wu et al., 2014). Interestingly, compared to WT control males, Syvn1 tended to be down-regulated in the HFD-fed GR-LKO males but upregulated in the AS GR-LKO males, resulting in a higher expression of Syvn1 in the GR-LKO AS males than GR-LKO AS females (Fig. 4M). Syvn1 is induced by ER stress (Wu et al., 2014). The higher SYVN1 expression in the GR-LKO AS males than in females may be due to elevated ER stress in these males. Glucocorticoid-induced leucin zipper (GILZ/TSC22D3) is a prototypical GR-target gene that plays a key role in mediating the anti-inflammatory response of GCs. Gilz was down-regulated by ethanol treatment in the WT AS mice (Fig. 4N). Surprisingly, Gilz tended to be further down-regulated in the GR LHet AS males but not in the GR-LKO AS males, suggesting that some compensatory pathway(s) may prevent the further down-regulation of GILZ in the GR-LKO AS mice. These data strongly support a key role of GR in protecting against fatty liver and liver injury in AS.

### Alteration of hepatic ERα signaling in GR-LKO AS mice

The lack of marked elevation of blood levels of ALT and bilirubin in GR LKO AS mice suggests that certain adaptive changes may help ameliorate severe liver injury in these mice. Studies of HFD plus acute binge drinking have shown that females are less prone to AS, likely due to sex differences in E2 signaling. Moreover, GR-LKO female AS mice were protected from liver injury. Thus, we explored changes in hepatic estrogen signaling. Hepatic ERα was female predominant and induced by GR deficiency and ethanol treatment in males (Fig. 5A). The GR-target gene SULT1E1 is a key enzyme for the inactivation of E2 (Gong et al., 2008). Hepatic expression of Sult1e1 was highly gene-dosage-dependent on GR in both genders (Fig. 5B). The ERα-target gene GDF15 maintains mitochondrial function and protects against NAFLD and ALD (Chung et al., 2017). We found gene-dosage-dependent induction of Gdf15 in the GR LHet and GR LKO AS mice (Fig. 5C). Loss of the ERα-target gene stress-responsive activating transcription factor 3 (ATF3) exacerbates liver damage via the deficiency of NRF2/heme oxygenase-1 and activation of mTOR/p70S6K/HIF-1α signaling pathways in inflammatory liver injury (Rao et al., 2015, Zhu et al., 2018b). Hepatic Atf3 expression was female-predominant and increased in the GR LHet and LKO AS mice (Fig. 5D). Hepatic induction of ATF3 may help ameliorate liver injury in these AS mice. Leptin deficiency enhances the sensitivity of rats to AH (Martínez-Uña et al., 2020). The ERα-target gene leptin receptor (Lepr) (Yin et al., 2017) was strongly induced in the female, but not male, WT AS mice (Fig. 5E, 5F). In contrast, Lepr was highly induced in both genders of GR-LKO AS mice. The active long isoform of the LEPR (Lepr-V1/Ob-RB) was similarly induced like the total Lepr, indicating that the functional Ob-RB accounts for the majority of Lepr mRNA induced in the GR LKO mice (Fig. 5E, 5F). Overexpression of the antioxidative enzyme quinone oxidoreductase 1 (NQO1) prevents alcohol-induced hepatic NAD+ depletion, thereby enhancing activities of NAD+-dependent enzymes and reversing alcohol-induced liver injury (Dong et al., 2021). Hepatic expression of Nqo1 in female mice was twice as high as that in male mice (Fig. 5G). Nqo1 expression remains unchanged in the WT AS mice but was induced in both genders of GR-LKO AS mice, suggesting that induction of the antioxidative enzyme NQO1 may help ameliorate oxidative liver injury in these mice. CYP2B family enzymes are important for the catabolism of unsaturated fatty acids (Damiri and Baldwin, 2018). Cyp2b9, a well-established female-specific ERα-target gene (Yamada et al., 2002), was highly induced in the HFD-fed and AS male GR LKO mice (Fig. 5H), which may help the clearance of unsaturated fatty acids in these mice. In summary, these data suggest that adaptive induction and activation of ERα and ERα-target genes may play a key role in protecting against liver injury in GR-LKO AS mice and early AS in humans.

**Figure 5.**
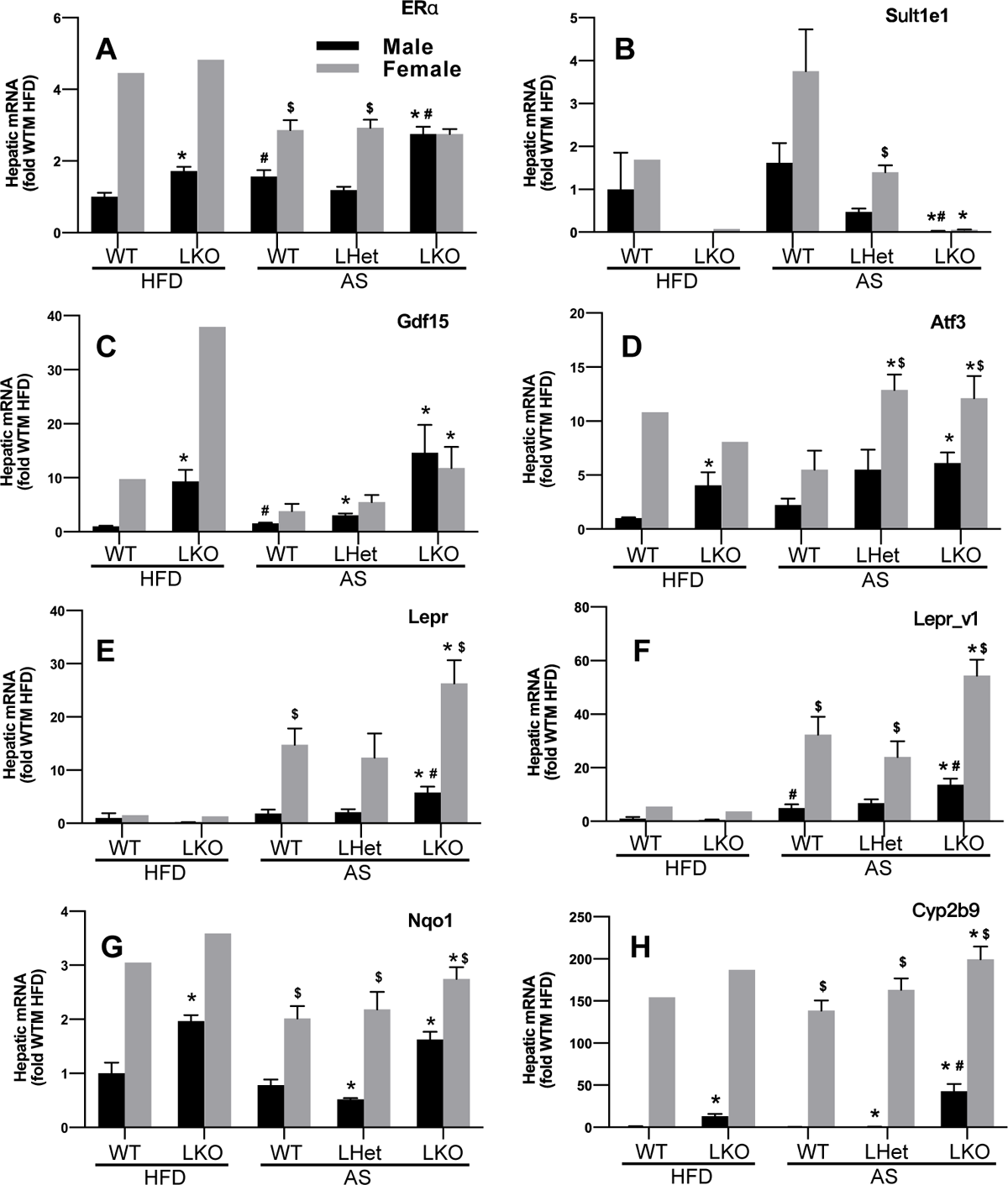
Hepatic mRNA expression of ERα and ERα-target genes in male and female wildtype (WT), GR liver-specific heterozygous (LHet) and GR liver-specific knockout (LKO) HFD-fed and alcoholic steatosis (AS) mice. Mice were fed a HFD for 3 weeks followed by ig administration of ethanol 5 g/kg (AS groups) or isocaloric maltose 9 g/kg (HFD groups) in the morning. All mice were sacrificed 9 h after ethanol treatment to collect livers for determination of mRNA by qPCR. N=3-7 per group, mean ± SE (pooled samples for WT-HFD and KO-HFD females). * p < 0.05 vs corresponding WT group; # p < 0.05 vs corresponding HFD group; $ p < 0.05 vs corresponding male group.

In contrast to the induction of ERα and ERα-target genes in GR LKO AS mice, our data mining found that human sAH had marked down-regulation of ERα (ESR1, ↓74%) and the ERα-target genes LEPR (↓80%), ATF3 (↓91%), and CYP2B6 (↓50%) (Dickmann and Isoherranen, 2013) (Fig. S1). Thus, down-regulation of hepatic ERα and ERα-target genes may play an important role in the pathogenesis of sAH in humans.

### Changes in hepatic protein expression in AS mice

As expected, hepatic nuclear (Fig. 6A male; Fig. 8A female) and cytosolic (Fig. 7A) GR proteins were gene-dosage-dependently decreased in GR LHet and LKO AS mice, demonstrating the highly efficient deletion of GR in hepatocytes in our inducible GR LHet and LKO mice. Hepatocyte nuclear factor-4 alpha (HNF4α) is a master regulator of liver development and function whose deficiency has been implicated to play a key role in the progression of AS in humans (Argemi et al., 2019). HNF4α controls liver metabolism and lipid homeostasis via crosstalk with diverse signaling pathways to regulate hepatic nutrient metabolism (Huang et al., 2020, Lu, 2016). Interestingly, HNF4α protein levels were strongly increased in the nucleus (Fig. 6B male; Fig. 8B female) and cytosol (Fig. 7B) of WT mice in this acute AS model, whereas the increase of nuclear HNF4α proteins by acute ethanol treatment was lost in both the male GR LHet and LKO AS mice, indicating a critical role of GR in maintaining hepatic protein levels of HNF4α in AS in males. In contrast, nuclear HNF4α proteins were unchanged in GR LHet females and only tended to be lower in the GR LKO females (Fig. 8B).

**Figure 6.**
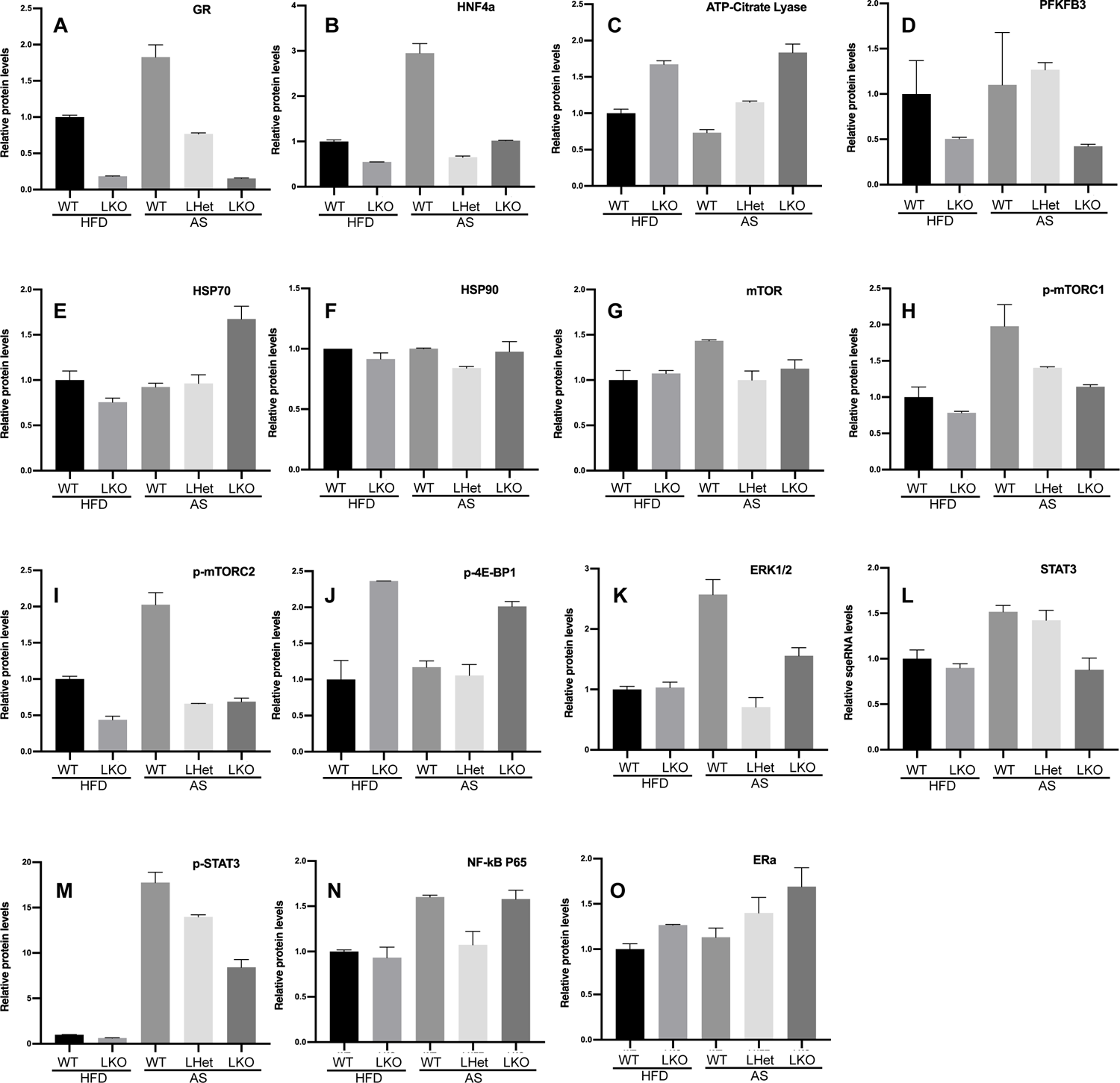
Quantification of Western blot of proteins in pooled liver nuclear extracts from male wildtype (WT), GR liver-specific heterozygous (LHet) and GR liver-specific knockout (LKO) HFD-fed and alcoholic steatosis (AS) mice. Mice were fed a HFD for 3 weeks followed by ig administration of ethanol 5 g/kg (AS groups) or isocaloric maltose 9 g/kg (HFD groups) in the morning. All mice were sacrificed 9 h after ethanol treatment to collect blood and tissues for analysis. Data were normalized to total proteins in each lane.

**Figure 7.**
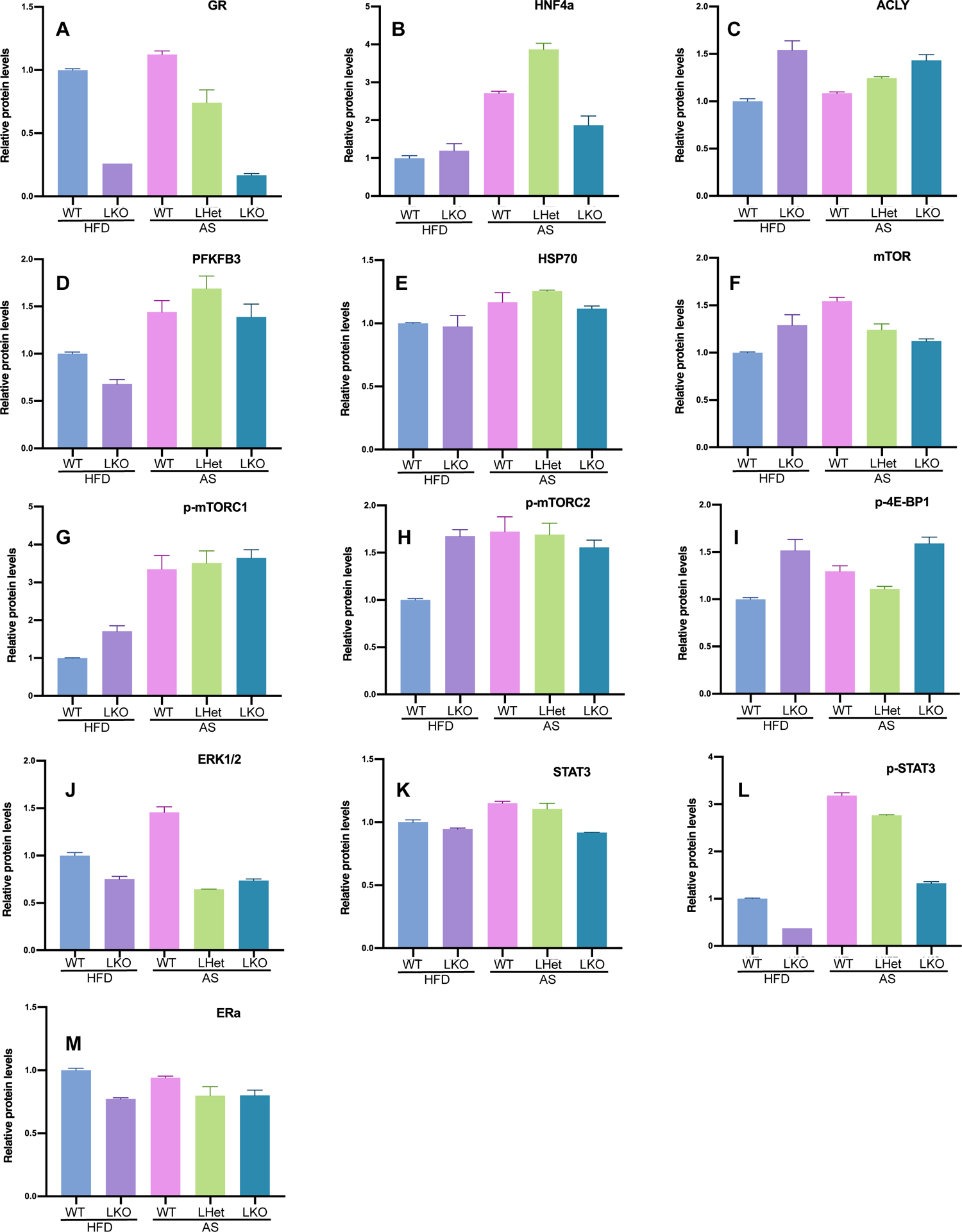
Quantification of Western blot of proteins in pooled liver cytosolic extracts from male wildtype (WT), GR liver-specific heterozygous (LHet) and GR liver-specific knockout (LKO) HFD-fed and alcoholic steatosis (AS) mice. Mice were fed a HFD for 3 weeks followed by ig administration of ethanol 5 g/kg (AS groups) or isocaloric maltose 9 g/kg (HFD groups) in the morning. All mice were sacrificed 9 h after ethanol treatment to collect blood and tissues for analysis. Data were normalized to total proteins in each lane.

**Figure 8.**
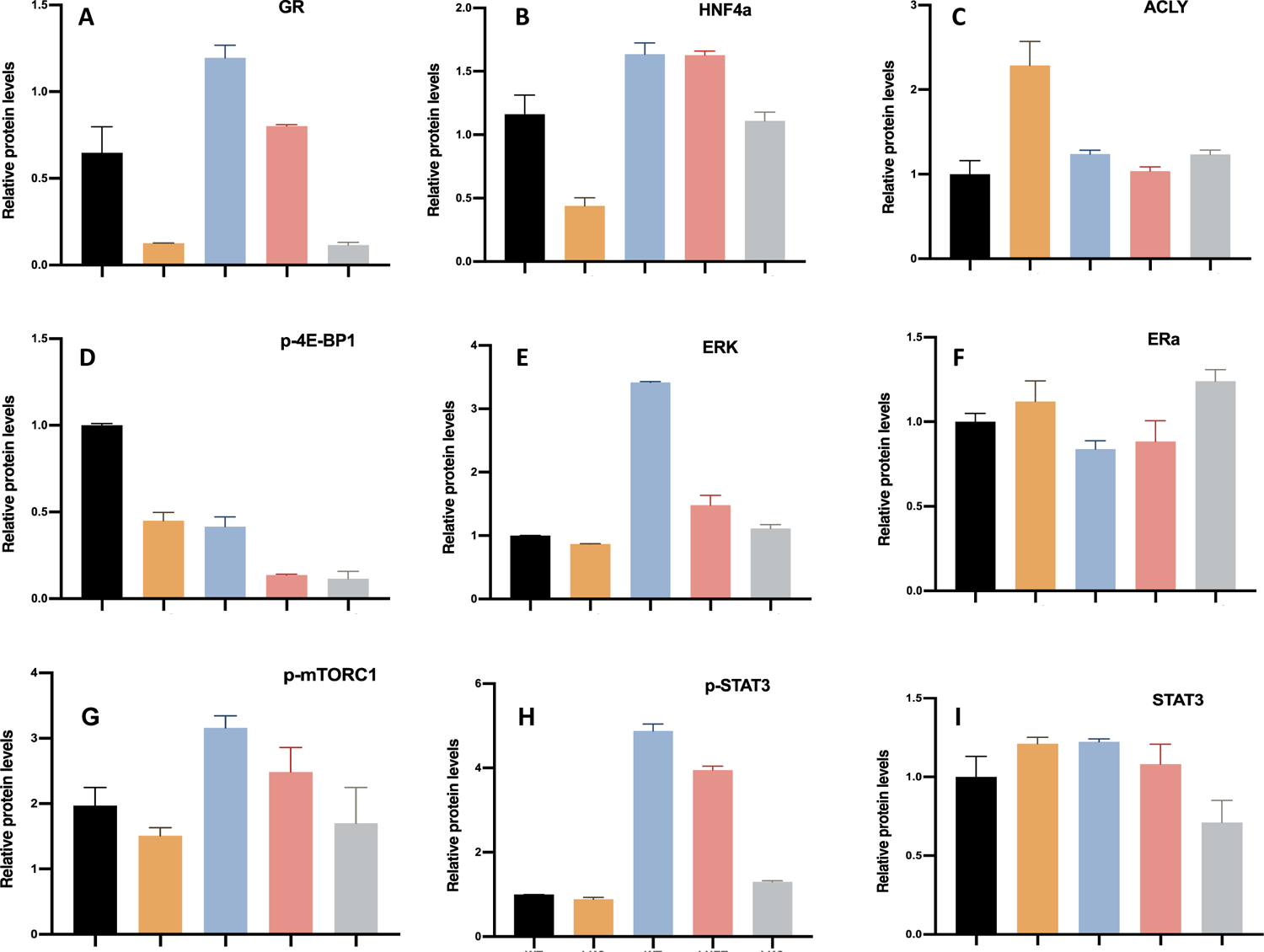
Quantification of Western blot of proteins in pooled liver nuclear extracts from female wildtype (WT), GR liver-specific heterozygous (LHet) and GR liver-specific knockout (LKO) HFD-fed and alcoholic steatosis (AS) mice. Mice were fed a HFD for 3 weeks followed by ig administration of ethanol 5 g/kg (AS groups) or isocaloric maltose 9 g/kg (HFD groups) in the morning. All mice were sacrificed 9 h after ethanol treatment to collect blood and tissues for analysis. Data were normalized to total proteins in each lane.

Consistent with the induction of Acly and down-regulation of Pfkfb3 mRNAs, nuclear levels of ACLY (Fig. 6C; Fig. 8C female) and PFKFB3 (Fig. 6D) proteins were elevated and decreased in HFD-fed and AS GR-LKO mice, respectively. Compared to the nucleus, cytosolic levels of ACLY proteins (Fig. 7C) increased to a less degree in the GR-LKO AS males, whereas cytosolic PFKFB3 proteins (Fig. 7D) were similarly elevated in the WT and GR-LKO AS males. An increase of heat shock protein 70 (HSP70) by E_2_ is hepatoprotective during ischemia-reperfusion injury (Shen et al., 2007). HSP70 proteins were increased in the nucleus (Fig. 6E), but not cytosol of GR-LKO AS mice (Fig. 7E), which may help ameliorate liver injury in the male GR-LKO AS mice. In contrast, hepatic expression of HSP90 remained unchanged in all the HFD-fed and AS mice (Fig. 6F).

Alcohol causes activation of mammalian target of rapamycin complex (mTOR), which increases lipogenesis and decreases autophagy (Chao et al., 2018). mTOR signaling complex 1 (mTORC1) is a key regulator of cell growth and nutrient signaling whereas mTORC1 regulates the actin cytoskeleton and activates Akt. mTORC1 contains mTOR phosphorylated predominantly on Ser2448 whereas mTORC2 contains mTOR phosphorylated predominantly on Ser2481 (Copp et al., 2009). As expected, phosphorylation of mTORC1 (p-mTORC1) and p-mTORC2 were increased in both the nucleus (Fig. 6G-I) and cytosol (Fig. 7G-I) of the WT male AS mice. Surprisingly, compared to male WT AS mice, p-mTORC1 and p-mTORC2 in the nucleus (Fig. 6G-I), but not the cytosol (Fig. 7G-I), of GR-LKO AS mice were decreased. The contribution of the selective inhibition of nuclear mTOR activation to the pathology in the GR-LKO AS mice warrants further investigation. mTOR promotes protein synthesis via phosphorylating the 70-kDa ribosomal protein S6 kinase (S6K1), and mTOR phosphorylates eIF4E-binding protein 1 (4E-BP1) on Thr-37 and Thr-46, resulting in inactivation of 4E-BP1 to allow cap-dependent translation to proceed (Gingras et al., 1999). Surprisingly, Thr-37 and Thr-46 p-4E-BP1 were increased in GR-LKO AS male mice (Fig. 6J nucleus; Fig. 7I cytosol) but decreased in the nucleus of females (Fig. 8D), despite decreased p-mTORC1 and p-mTORC2 in both genders. In the nucleus, 4E-BP1 binds to and retains eIF4E in the nucleus and thus affects the nuclear-cytoplasmic transport of certain mRNAs (Rong et al., 2008). Interestingly, 4E-BP1 levels are decreased in male, but not female livers of aging mice (Tsai et al., 2016). Overexpression of 4E-BP1 protects males, but not females, against hepatosteatosis in HFD-fed mice (Tsai et al., 2016). In contrast, knockout of S6K1 only prolongs the lifespan of female mice (Selman et al., 2009). Currently, the mechanism of the gender-difference in hepatic phosphorylation of 4E-BP1 in GR LKO AS mice is unclear. In addition to mTORC1/C2, 4E-BP1 can also be phosphorylated by other kinases such as glycogen synthase kinase 3β and p38 (Qin et al., 2016). Thus, it is likely that kinase(s) other than mTORC1/C2 may contribute to the aberrant increased phosphorylation of 4E-4P1 in male GR-LKO AS mice.

Activation of extracellular signal-regulated kinase (ERK) is associated with liver injury in AH (Aroor et al., 2011). In the GR-LKO AS mice, nuclear (Fig. 6K male; Fig. 8E female) and cytosolic (Fig. 7J) p-ERK was increased to less degrees than in the WT AS mice, which indicates that GR regulates hepatic activation of MAPK/ERK signaling pathway. Activation of ERK ameliorates liver steatosis in leptin receptor–deficient (db/db) mice via stimulating ATG7-dependent autophagy (Xiao et al., 2016). Loss of both ERK1 and ERK2 in hepatocytes causes spontaneous liver injury in mice (Cingolani et al., 2022). Likewise, activation of signal transducer and activator of transcription 3 (STAT3) by interleukin-6 protects against HFD-fed-induced fatty liver and AS (Wang et al., 2011). Hepatic protein expression of STAT3 was moderately increased (Fig. 6L nucleus; 7K cytosol), whereas p-STAT3, the active form, was dramatically increased by ethanol treatment in the nucleus (Fig. 6M) and cytosol (Fig. 7L) of WT AS mice. GR deficiency attenuated the increase of hepatic STAT3 phosphorylation by AH, indicating less liver protective effects by STAT3 in the GR-LKO AS mice. After activation by tumor necrosis factor, nuclear factor kappa B (NF-kB) translocates to the nucleus to promote cytokine production and cell survival. Nuclear NF-kB levels were similarly increased moderately in the WT and GR-LKO AS mice (Fig. 6N). This is consistent with the previous study that GR inhibits the transcriptional activity of NF-kB by disturbing the interaction of p65 with the basal transcription machinery without affecting the protein levels of p65 (De Bosscher et al., 2000).

Consistent with the mRNA results (Fig. 5A), nuclear ERα proteins increased in male AS mice (Fig. 6O), but the change was less prominent in females (Fig. 8F). This may be due to higher basal hepatic expression levels of ERα in females than in males.

### Effect of GR-LKI on AH in middle-aged mice

Ten days of chronic drinking (6.7% ethanol in liquid diet, F1258SP, Bio-Serv) plus one binge alcohol (5g/kg) caused marked macrovesicular steatosis in WT AH mice. WT mice fed a control liquid diet (F1259SP, Bio-Serv) and the GR-LKI AH mice had only mild steatosis (Fig. 9A). sAH is frequently associated with liver fibrosis in humans. Trichrome staining showed perisinusoidal fibrosis in WT AH mice, but not in WT control or GR-LKI AH mice (Fig. 9B). Consistent with marked steatosis in WT AH mice, liver/body weight ratio (Fig. 9C) and hepatic TG levels (Fig. 9D) increased 29% and 105% in WT AH mice, but were unchanged in GR-LKI AH mice. Blood levels of TG tended to increase (↑30%), whereas cholesterol tended to decrease (↓40%) in WT AH mice (Fig. 9E). In contrast, both TG and cholesterol tended to be lower in GR-LKI AH mice than GR-LKI control mice. Blood ALT tripled in WT AH mice, indicating significant liver injury, whereas ALT was unchanged in GR-LKI AH mice (Fig. 9F). Liver-specific loss of the lipogenic Perilipin 2 (Plin2) alleviates diet-induced hepatic steatosis, inflammation, and fibrosis (Najt et al., 2016). Our qPCR data (Fig. 9G) showed that mRNA expression of key lipogenic genes Scd1 and G0s2 tended to be down-regulated more markedly in GR-LKI AH mice than WT AH mice, whereas Plin2 tended to be induced in WT AH mice but unchanged in GR-LKI AH mice, resulting in significantly lower Plin2 in GR-LKI AH mice (Fig. 9G). Proinflammatory cytokines Cxcl1 and lipocalin 2 (Lcn2), important contributors to AH (Asimakopoulou et al., 2016, Chang et al., 2015, Chen et al., 2020), were strongly induced in both WT and GR-LKI AH mice (Fig. 9H). Plasminogen activator inhibitor-1 (PAI-1) plays a key role in steatosis, inflammation and fibrosis (Ghosh and Vaughan, 2012, Levine et al., 2021, Beier and Arteel, 2012). PAI-1 was markedly induced 22-fold in WT AH mice, but only induced 4-fold in GR-LKI AH mice, which was associated with 60% lower collagen 1a1 (Col1a1) in GR-LKI AH mice (Fig. 9I). Hepatic total GR mRNAs tended to increase 4-5 fold in the GR-LKI mice than WT mice (Fig. 9J). Interestingly, Sult1e1 was the only tested GR-target gene significantly altered in GR-LKI control mice, and Sult1e1 was induced to a much higher level in GR-LKI AH mice than WT AH mice (Fig. 9J).

**Figure 9.**
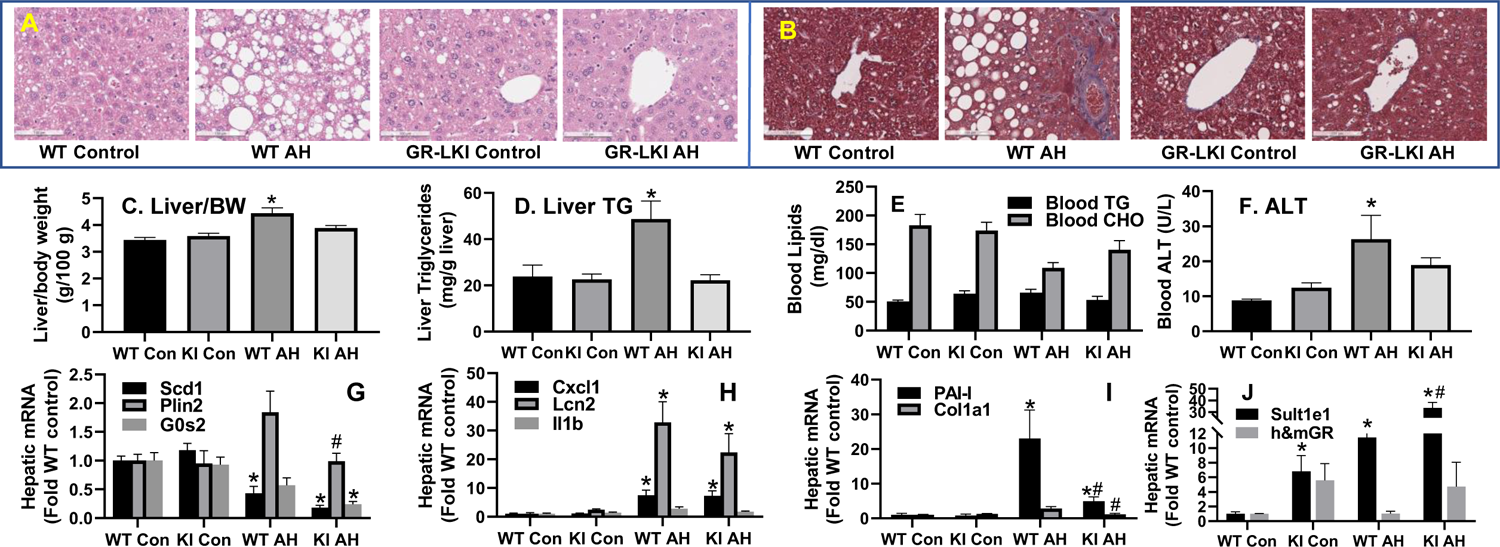
Liver histology (A: H & E; B: Trichrome staining), liver/body weight (BW) ratio (C), hepatic triglycerides (D), blood lipids (E), blood ALT (F), and hepatic mRNA expression (G-J) determined by qPCR (normalized to peroxiredoxin 1) in male middle-aged (10-12 month-old) mice with liver-specific knockin of GR (GR-LKI) and age-matched wildtype (WT) littermate mice with chronic-plus-binge (E10+1B)-induced alcoholic hepatitis (AH). N=4-7 per group, Mean ± SE. * p<0.05 vs WT control (WT Con); # p<0.05 vs WT AH.

## DISCUSSION

In the present study, we found that hepatic GR deficiency worsened steatosis and liver dysfunction in both genders of HFD-fed mice and AS mice, but only increased the marker of liver injury in male AS mice. Certain key GR-target genes important for cytoprotection and lipid metabolism were deregulated in GR LKO AS mice. In contrast, LKI of GR protects against liver injury and steatohepatitis in chronic-plus-binge-induced AH in male middle-aged mice. For the first time, our data identified a novel important role of hepatic GR in protecting against AS and AH. Interestingly, hepatic expression of ERα was induced, and the key E2-inactivating enzyme SULT1E1 was markedly down-regulated in GR knockout AS mice, suggesting enhanced E2-ERα signaling in these mice. Multiple ERα-target cytoprotective genes were induced in GR LKO AS mice, which may help ameliorate liver injury in these mice. The findings of this study will help us determine the mechanistic role of GR in alcoholic and non-alcoholic fatty liver disease and develop targeted drug therapies to treat alcoholic/non-alcoholic steatohepatitis.

The present study provides the first evidence of an imperative role of hepatocellular GR in the protection against alcoholic steatosis. Certain key GR-target genes important for cytoprotection and lipid catabolism were gene-dosage-dependently down-regulated in the GR LHet and GR LKO mice. Particularly, GR was found to have a critical role in maintaining hepatic mRNA expression and serum levels of albumin. Additionally, consistent with a previous report (Quagliarini et al., 2019), the present study identified an essential role of hepatic GR in the protection against HFD-induced NAFLD. Albeit traditionally viewed as different diseases, ALD and NAFLD share many similarities regarding the etiology and pathophysiology (Idalsoaga et al., 2020). The vital roles of hepatic GR in the protection against both the ALD and NAFLD strongly support the current usage of glucocorticoids as the first-line drugs to treat sAH and the potential usage of liver-targeting GCs to treat moderate AH and NAFLD as well.

Changes in GR levels at different stages in AH are also a point of interest. GR appears to be activated in the liver of WT AS mice (Fig. 6A, 8A) despite the lack of increase in the circulating corticosteroid (data not shown). In this regard, ethanol treatment has been shown to induce the GR-target gene GILZ in the cultured human lung epithelial cells via increasing nuclear translocation of GR (Gomez et al., 2010). Thus, hepatic GR may be activated by ethanol or its metabolites. Despite activation of GR, hepatic expression of the prototypical GR-target gene GILZ tended to be down-regulated in the WT AS mice. It has been shown that the induction of GILZ by GR requires the transcription factor FOXO3 (Asselin-Labat et al., 2005). We found that Foxo3 mRNA was down-regulated by ethanol in the WT AS mice (data not shown), which may contribute to the trend of down-regulation of GILZ in WT AS mice. In contrast to the activation of GR in acute AS, hepatic expression of GR and GR-target genes are markedly down-regulated in sAH patients (Lu, 2022). Further study on the mechanism of hepatic GR deficiency in sAH is warranted.

The present study demonstrates important roles of GR in the activation of hepatic STAT3 and maintenance of HNF4α protein levels in AS. Activation of STAT3 by IL-6 protects against HFD-fed-induced fatty liver and AS (Wang et al., 2011). GR induces IL-6 receptor (IL-6R) in hepatocytes (Snyers et al., 1990). We found down-regulation of IL-6R in GR-LKO AS mice (data not shown), which may be the underlying mechanism of impaired STAT3 activation in these mice. GR and STAT3 crosstalk extensively to regulate gene expression. Particularly, GR and STAT3 synergize to induce Mt1 (Langlais et al., 2012). Consistently, the dramatic induction of Mt1 in WT AS mice (Fig. 4C) is associated with activation of GR and STAT3 in these mice, whereas the loss of Mt1 induction in the GR LKO AS mice is associated with attenuated activation of STAT3. In parallel, the human ortholog MT1X is dramatically down-regulated in human sAH (Lu, 2022). Impaired liver regeneration is a major contributor to liver failure and mortality in sAH (Dubuquoy et al., 2015). Metallothioneins as well as hepatocellular GR and STAT3 are important for liver regeneration (Cherian and Kang, 2006, Shteyer et al., 2004, Wang et al., 2011). The role of hepatic GR-STAT3-MT in liver protection and regeneration warrants further investigation. In addition to a critical role in maintaining hepatic metabolic homeostasis, HNF4α is also required for liver regeneration by promoting the differentiation of mature hepatocytes (Huck et al., 2019). Deficiency of HNF4α plays a central role in the pathogenesis of sAH (Argemi et al., 2019). The important roles of GR in maintaining hepatic STAT3 and HNF4α in AH, key factors for hepatocyte proliferation and differentiation, suggest that deficiency of hepatocellular GR will impair liver regeneration in sAH.

Despite the severe steatosis and liver dysfunction developed in the GR-LKO AS mice, the moderate increase of ALT and lack of hyperbilirubinemia is distinct from those in the patients with sAH. It is well known that many severe liver diseases require a second hit. Studies of HFD plus acute binge drinking have shown that females are less prone to AH, likely due to sex differences in E2 signaling (Gao et al., 2017). Hepatic ERα protects against fatty liver disease and promotes liver regeneration (Zhu et al., 2013, Zhu et al., 2018a, Della Torre et al., 2011, Lee et al., 2019, Uebi et al., 2015, Tsugawa et al., 2019, Dubuquoy et al., 2015). In the early stage of AS, hepatic ERα increases, which may help ameliorate AS. E2 is inactivated by hepatic SULT1E1. GR inhibits ERα via physical interaction with ERα and induction of SULT1E1 (Gong et al., 2008, Karmakar et al., 2013, West et al., 2016, Bolt et al., 2013). The present study discovered a critical role of GR in maintaining hepatic SULT1E1 expression in AS. We found certain adaptive changes in the GR-LKO AS mice that may help ameliorate liver injury in these mice, among which the induction of ERα and ERα-target genes such as LEPR, ATF3, GDF15, NQO1, and CYP2B9 was the most striking changes adaptive to GR deficiency in these mice. These data suggest that in the early stages of AH, to protect the liver cells, GR and ERα levels increase. E2/ERα signaling inhibits hepatic lipogenesis (Qiu et al., 2017b). Importantly, hepatic ERα levels correlate negatively with ALD severity (Colantoni et al., 2002, Eagon et al., 2001, Villa et al., 1988). E2 is largely decreased after menopause, which occurs earlier in alcoholic women (Mello, 1988, Becker et al., 1991), and menopause increases the risks of NAFLD and ALD (DiStefano, 2020). ERα also regulates lipid metabolism in males (Qiu et al., 2017a, Della Torre et al., 2016). ERα has recently been recognized as a relevant molecular target for the prevention of NAFLD (Della Torre, 2020). However, its role in AH remains poorly understood. ERα levels are elevated in hepatitis patients who abuse alcohol (Bell et al., 1995). In patients with sAH, hepatic ERα levels drop dramatically (Vandegrift et al., 2020, McKetta and Keyes, 2019). This suggests that elevated ERα levels in the early stage of AS may help ameliorate liver injury. In contrast, down-regulation of ERα in the late stage of AH, particularly in postmenopausal women, may play an important role in the pathogenesis of sAH in these patients. The role of hepatocellular ERα in the protection against liver injury in the GR-LKO AS mice, aging AS mice, and sAH patients warrants further investigation.

The present study demonstrates a critical role of hepatic GR in regulating SULT1E1 expression in alcoholic liver disease. Hepatic expression of SULT1E1 is downregulated in patients with sAH (Fig. S1) and cirrhosis (Yalcin et al., 2013). In addition to its high-affinity substrate estrogen, SULT1E1 can also sulfate other compounds, namely dehydroepiandrosterone (DHEA), pregnenolone, and thyroid hormone (Yi et al., 2021). SULT1E1 deficiency worsens liver ischemia/reperfusion injury and increases mortality during polymicrobial sepsis in male mice (Guo et al., 2015, Chai et al., 2015). Currently, little is known about the role of SULT1E1 in alcoholic liver disease. We found that the downregulation of SULT1E1 is associated with induction of ERα-target genes and mild liver injury in GR LKO AS mice, and induction of SULT1E1 in GR LKI AH mice is associated with protection against liver injury and steatohepatitis. The roles of GR-SULT1E1 pathway in steroid hormone metabolism, AH pathogenesis, and susceptibility of AH patients to sepsis warrant further investigation.

Currently, there is no conclusion regarding gender difference in binge-drinking-induced liver dysfunction (Wilsnack et al., 2018). In the present study, eight-week-old mice were fed HFD for 3 weeks, followed by binge alcohol to induce acute AS in these mice. Consistent with literature (Gao et al., 2017, Spruiell et al., 2015, Li et al., 2017), we found less liver injury in the AS females than AS males. Interestingly, in a large population study, compared to non-binge drinkers, binge drinking increases the odd of liver steatosis in the obese men but not the obese women (Lau et al., 2015). In contrast, women are considered more susceptible to chronic drinking-induced ALD (Delacote et al., 2020, Buzzetti et al., 2017, Ballestri et al., 2017, Mauvais-Jarvis et al., 2020, Li et al., 2019), partly due to lower gut alcohol clearance and higher gut permeability to endotoxins (Eagon, 2010). In the mouse model of chronic AH induced by chronic plus binge ethanol feeding, females are either similar or more susceptible to liver injury than the males (Bertola et al., 2013a). Plasma endotoxin levels are higher in ovariectomized rats given estrogen replacement than in ovariectomized animals exposed chronically to ethanol (Yin et al., 2000). Thus, estrogens may protect the liver per se from injury via activating the hepatocellular ERα but increases gut permeability to endotoxin or endotoxin production during chronic drinking, resulting in ultimate either similar or aggravated liver injury in females than males in the chronic plus binge-induced AH. Future study on the role of hepatic GR and ERα in gender-divergent AH pathogenesis in the chronic plus binge model is warranted.

Aging humans and rodents have markedly increased alcoholic liver injury and decreased liver regeneration (Ramirez et al., 2017). The majority of sAH patients are middle-aged (mean age of 53 years) (Dugum and McCullough, 2016). Mice reach postmenopausal at 12 months of age (Diaz Brinton, 2012). Thus, 12-month-old mouse liver will be similar to middle-aged human liver. In this study, we found that in 12-month-old male WT mice, chronic plus binge (E10d+1B) induced marked steatohepatitis and significant fibrosis, which is similar to human AH. Liver-specific expression of the human GR mRNA in GR LKI mice markedly protected against liver injury, steatosis, and fibrosis, without significant effect on hepatic induction of inflammatory cytokines. PAI-1 plays a key role in steatosis, inflammation fibrosis, and thrombosis (Ghosh and Vaughan, 2012, Levine et al., 2021, Beier and Arteel, 2012). GCs strongly induce PAI-1 in adipose tissue but down-regulate PAI-1 in mouse liver (Tamura et al., 2015). Consistently, we found strong down-regulation of PAI-1 in GR LKI AH mice, which is associated with markedly ameliorated steatosis and fibrosis in these mice. Additionally, we found that GCs strongly down-regulated PAI-1 in lipopolysaccharides-treated primary human hepatocytes (manuscript in revision). In both humans and mice, blood levels of PAI-1 correlate with liver steatosis and hepatic PAI-1 mRNA expression (Alessi et al., 2003, Zheng et al., 2020). The role of hepatic GR-PAI-1 pathway in AH pathogenesis and the potential of serum PAI-1 as biomarker of GC therapy in AH warrant further investigation.

In conclusion, using models of alcohol-induced liver steatosis and AH in mice with GR LKO and LKI, the present study provides strong evidence that hepatic GR deficiency plays a crucial role in the pathogenesis of alcoholic steatosis induced by HFD plus binge, and liver-specific overexpression/activation of GR protects against chronic-plus-binge-induced AH in middle-aged mice. These novel findings, in contrast to the known detrimental effects of activation of intestinal and adipose GR on steatohepatitis (Shukla et al., 2022, Shen et al., 2017), strongly support the development of liver-specific GR activation as an novel approach to improve sAH treatment.

## ACKNOWLEDGEMENTS

We would like to thank Dr. Pierre Chambon (IGBMC, Illkirch, France) for providing the SA^+/CreERT2^ mice (Schuler et al., 2004).

## AUTHOR CONTRIBUTIONS

YW: Conducted experiments, analyzed data, and wrote the draft manuscript; HL: Obtained grant funding, developed, and designed research project goals and experiments, conducted experiments, analyzed data, outlined, revised, and conducted final edition of manuscript.

