## Supplemental figure for "Liver-specific glucocorticoid action in alcoholic liver disease: study of glucocorticoid receptor knockout and knockin mice"

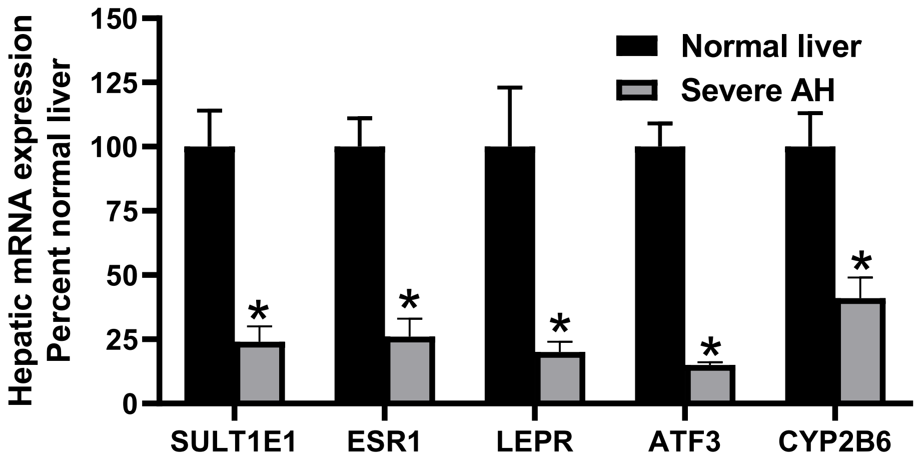


Supplemental Figure 1. Data mining of microarray analysis (GSE28619) of hepatic mRNAs in humans with severe alcoholic hepatitis (AH) (normalized to β-actin). N = 7-15 per group, mean ± SE. * $p < 0.05$ verus normal livers.

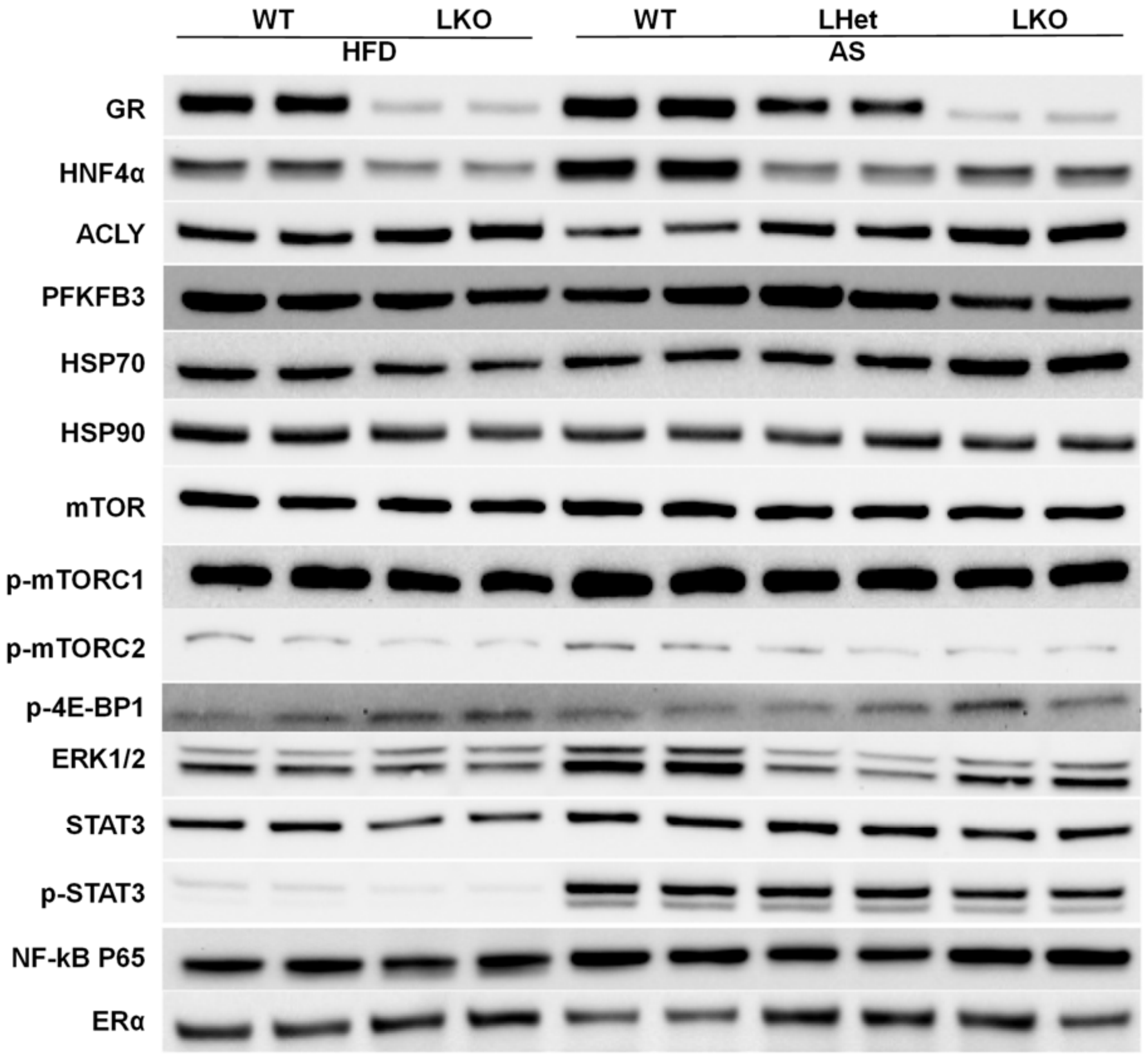


Supplemental Figure 2. Western blot determination of proteins in pooled liver nuclear extracts from male wildtype (WT), GR liver-specific heterozygote (LHet) and GR liver-specific knockout (LKO) HFD-fed and alcoholic steatosis (AS) mice. Mice were fed a HFD for 3 weeks followed by ig administration of ethanol 5 g/kg (AS groups) or isocaloric maltose 9 g/kg (HFD groups) in the morning. All mice were sacrificed 9 h after ethanol treatment to collect blood and tissues for analysis. Pooled samples were run in duplicates.

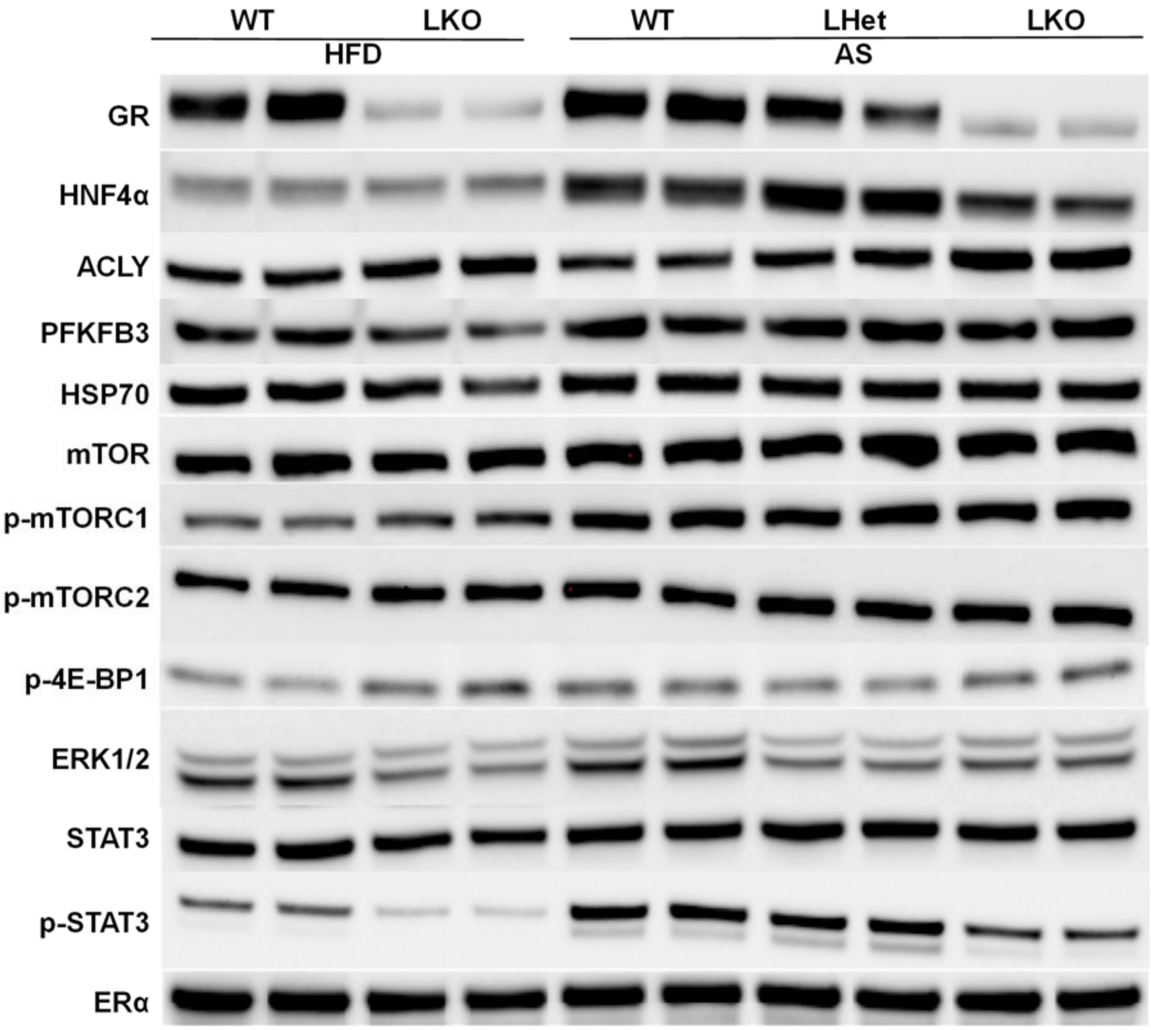


Supplemental Figure 3. Western blot determination of proteins in pooled liver cytosolic extracts from male wildtype (WT), GR liver-specific heterozygote (LHet) and GR liver-specific knockout (LKO) HFD-fed and alcoholic steatosis (AS) mice. Mice were fed a HFD for 3 weeks followed by ig administration of ethanol 5 g/kg (AS groups) or isocaloric maltose 9 g/kg (HFD groups) in the morning. All mice were sacrificed 9 h after ethanol treatment to collect blood and tissues for analysis. Pooled samples were run in duplicates.

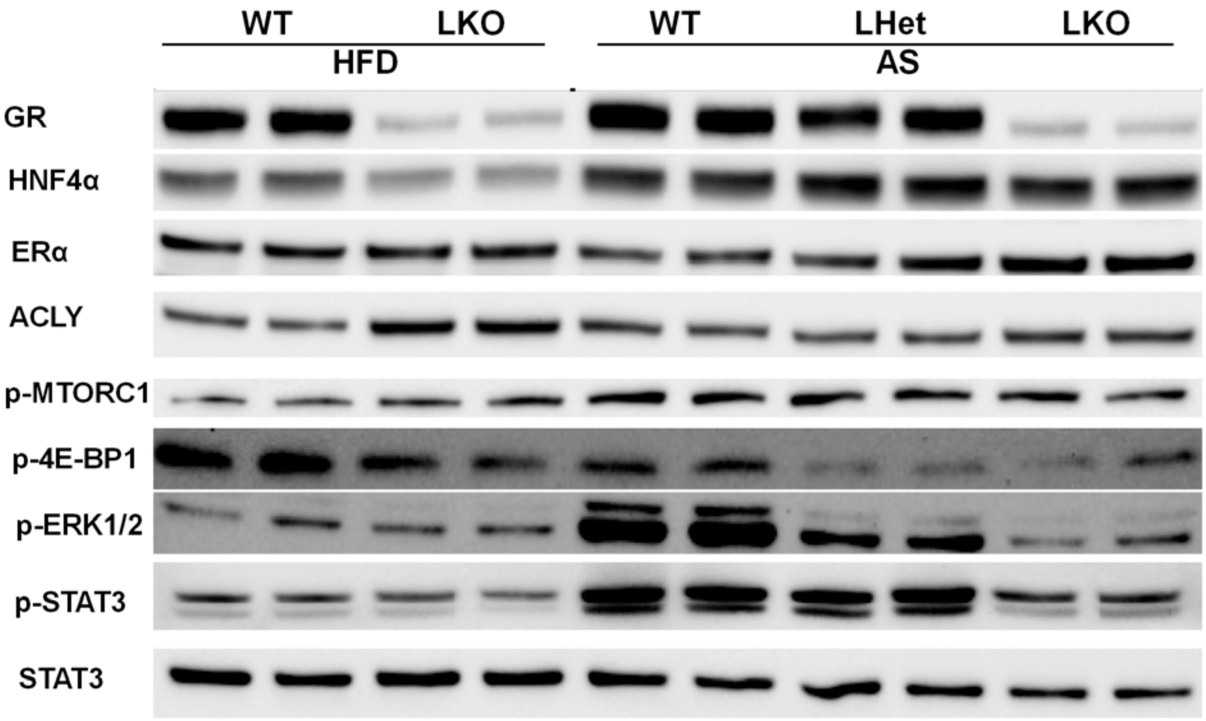


Supplemental Figure 4. Western blot determination of proteins in pooled liver nuclear extracts from female wildtype (WT), GR liver-specific heterozygote (LHet) and GR liver-specific knockout (LKO) HFD-fed and alcoholic steatosis (AS) mice. Mice were fed a HFD for 3 weeks followed by ig administration of ethanol 5 g/kg (AS groups) or isocaloric maltose 9 g/kg (HFD groups) in the morning. All mice were sacrificed 9 h after ethanol treatment to collect blood and tissues for analysis. Pooled samples were run in duplicates.
