## Supplemental Table 1 for "Liver-specific glucocorticoid action in alcoholic liver disease: study of glucocorticoid receptor knockout and knockin mice"

Supplemental Table 1. List of primers used in qPCR.

| | | |
|---|---|---|
| Acly | CCCAGTGAACAACAGACCTAT | ACAAAGATGGTGACCTCATGC |
| Alb | TCCGTCAGAGAATGAAGTGCT | GCAGTTTGCTGGAGATAGTCG |
| Apoa4 | GGGCTGAGGTCACTTCGGA | ATCCCCAAGTTTGTCCTGGAA |
| Atf3 | CTTCCCCAGTGGAGCCAATC | GACCTGGCCTGGATGTTGAA |
| CXCL10 | GTCATTTTCTGCCTCATCCTG | CATCGTGGCAATGATCTCAACA |
| Cyp2b9 | CAGTGCTCCACGAGACTACAT | AAGAGTTGGTAGCCGGTGTG |
| Cyp39a1 | CCACAGGACTGTTTTAGAAAGC | CAGATTCCAGAATGCACCACT |
| Cyp7b1 | CGGAAATCTTCGATGCTCCAA | AATCGGGGTGCTGAATACCTAA |
| Esr1 | CTTTAAGAGAAGCATTCAAGGAC | TCATGCCCACTTCGTAACACT |
| Gdf15 | CATGCCAACCAGAGCCGAGA | GGTTGACGCGGAGTAGCAG |
| Lepr | ATGGATGTAAAAGTTCCTATGAG | TTTCCCACATCTTCTGACCAC |
| Lepr_v1 | TGTCCTACTGCTCGGAACAC | TTCTGAAATGGGTTCAGGCTCC |
| Lpin1 | TGTTCAAGAGACTGACAACGAT | AGGCACCTGATTCTGTCTACA |
| Mt1 | CCCAACTGCTCCTGCTCCA | GGTAGAAAACGGGGGTTTAGT |
| Nqo1 | TGAGCTGAAGGACTCGAAGAA | CCACTGCAATGGGAACTGAAA |
| Nr3c1 | GAGCTAGGAAAAGCCATTGTC | TAAGGAGATTTTCAACCACATCAT |
| Pgk1 | CCCAGAAGTCGAGAATGCCT | CTCGGTGTGCAGTCCCAAAA |
| Scd1 | CTACGACAAGAACATTCAATCC | AGAAGCCCAAAGCTCAGCTAC |
| Srebp1c | ACGGAGCCATGGATTGCACA | CTGTCTCACCCCCAGCATAG |
| Sult1e1 | TCCGTATGGTTCCTGGTATGA | GTTGAACGATTCTGTCCACAAG |
| Tsc22d3 | CACCCTTGGAGTCACTTCTCT | CCGCTATAGGATAGGCTTTGG |
| Ubd | ATGGCTTCTGTCCGCACCTG | TTTTGGAGTCTAGCAGAAGGAT |
